# The L-lactate dehydrogenase LldD contributes to oxidative stress resistance, survival from neutrophils, and host colonization in *Neisseria gonorrhoeae*

**DOI:** 10.1101/2025.11.17.688940

**Authors:** Jerri M Lankford, Willis E Barr, Cole A Andersen, Amitha A Karuppiah, Keena S Thomas, Ian J Glomski, Alison K Criss, Aimee D Potter

**Affiliations:** Department of Microbiology and Immunology, University of Iowa Carver College of Medicine, Iowa City, Iowa, USA; Department of Microbiology, Immunology, and Cancer Biology, University of Virginia, Charlottesville, Virginia, USA

**Author notes:** Address correspondence to Aimee D. Potter,. **Competing interests:** The authors have declared that no competing interests exist.

**Keywords:** Neisseria gonorrhoeae, lactate dehydrogenase, oxidative stress resistance, neutrophils, host-pathogen interactions, bacterial metabolism

## Abstract

Metabolic adaptation to the host environment is a key determinant of bacterial pathogenesis, enabling both colonization and invasive disease. This is particularly true for *Neisseria gonorrhoeae* (Gc), the causative agent of gonorrhea, which lacks effector-injecting secretion systems or toxins. Gc infection triggers a rapid influx of neutrophils (PMNs) that typically kill bacteria through multiple mechanisms, including a potent oxidative burst. Despite this, Gc exhibits remarkable resistance to reactive oxygen species and readily replicates in the presence of PMNs, which is in part due to the consumption of PMN-derived lactate. Previous studies demonstrated that the lactate permease, LctP, is required for oxidative stress resistance in Gc and host colonization in a murine model of gonorrhea, suggesting that lactate utilization contributes to virulence. Gc encodes four lactate dehydrogenases (LDHs) with distinct regulation and mechanisms, including two L-LDHs, LldD and LutACB. Although either enzyme alone supports L-lactate utilization, we found that both are required for full fitness during co-colonization with PMNs, indicating some non-redundant roles. Furthermore, LldD, but not LutACB, enhances oxidative stress resistance and is required for Gc colonization in a murine model of gonorrhea, whereas LutACB is dispensable. These findings identify LldD as a key factor promoting oxidative stress resistance, survival during PMN challenge, and host colonization.

## Introduction

*Neisseria gonorrhoeae* (the gonococcus, Gc) causes the sexually transmitted infection gonorrhea. Gonorrhea is frequently asymptomatic and can lead to severe clinical consequences if left untreated, especially in women, including pelvic inflammatory disease, ectopic pregnancy, and infertility (1–3). The success of Gc as a pathogen reflects its remarkable capacity to adapt to the host environment, particularly the nutrient and immune landscape of the genital tract (4–9). Lactate, a major metabolic end product of microbiome constituents and mammalian cells within the female genital tract, represents one of the most abundant metabolites Gc is exposed to during infection (10, 11). Lactate concentrations within the genital tract are widely estimated to be around 6mM; however, measurements as high as 20mM or even 111mM have been reported in women with *Lactobacillus* dominated microbiomes (12–17). Within this niche, the distribution of L-lactate and D-lactate isomers is roughly balanced but favors D-lactate (∼55%) (18). Because Gc is metabolically restricted to consumption of glucose, L-lactate, D-lactate, and pyruvate as primary carbon sources, lactate is a critical nutrient to support colonization and virulence (19).

Gc can consume both L- and D-lactate by the lactate permease, LctP. In the presence of glucose, lactate contributes directly to energy production, accelerates the emergence from lag phase, and promotes a 20% increase in logarithmic growth rate (10–12, 20, 21). The stimulation of Gc metabolism by lactate is thought to support Gc virulence by enhancing replication rate within the host and enhancing synthesis of proteins, pentose precursors, and lipooligosaccharide (20, 21). Mutants lacking LctP display increased sensitivity to complement-mediated killing due to reduced surface modification of lipooligosaccharide with sialic acid present in serum (i.e. sialylation), which is supported by lactate-stimulated metabolic activity (12, 22). Sialylated Gc are less susceptible to killing by primary human neutrophils (polymorphonuclear cells, PMNs) and elicit a less potent oxidative burst (23). Although PMN-derived reactive oxygen species (ROS) are not thought to kill Gc directly, LctP also promotes resistance to H_2_O_2_ mediated killing (24). Together, these mechanisms are thought to promote Gc survival *in vivo*. In support of this hypothesis, LctP is required for bacterial fitness in a mouse vaginal infection model of gonorrhea (12). Based on these data, it is apparent that the lactate permease serves as a virulence determinant in Gc, presumably due to its role in lactate transport. Potentially confounding these previous reports, however, LctP was later found to transport pyruvate in addition to D- and L-lactate (25).

To unambiguously address the role of lactate in Gc pathogenesis, we have turned to interrogating the Gc enzymes responsible for L-lactate catabolism. Gc encodes four known enzymes that catalyze the oxidation of lactate to pyruvate: The L-lactate dehydrogenases (L-LDHs) LldD and LutACB, and the D-lactate dehydrogenases (D-LDHs) LdhD and LdhA (26, 27). LldD, LutACB, and LdhD are quinone-dependent LDHs, and support Gc respiration by shuttling electrons from lactate to the electron transport chain (26). LdhA is the sole NAD^+^-dependent LDH in Gc and is thought to primarily contribute to D-lactate production, rather than consumption (26). Previous work has shown that *ldhD* and *ldhA* mutants exhibit a survival defect in the presence of PMNs (26). Although no defect was observed for an *lldD* mutant, the role of LutACB, which was uncharacterized prior to 2019, was not interrogated (27).

Furthermore, none of the LDHs have been investigated for their contribution to oxidative stress, or in murine models of gonorrhea. Our understanding of how Gc lactate metabolism contributes to virulence is therefore incomplete. Here, we investigate how use of L-lactate by the L-LDHs, LldD and LutACB, impacts survival in key models of Gc infection *in vitro* and *in vivo*.

## Results

### LldD and LutACB are both required for L-Lactate utilization

LldD and LutACB were previously identified as the sole L-LDHs in Gc (27). To confirm this was true in our strain background, we generated null mutants of *lldD* (Δ*lldD*) and *lutACB* (Δ*lutACB*) in a derivative of Gc strain FA1090 lacking opacity associated (Opa) proteins (deemed Opaless) (28). As expected, both Δ*lldD* and Δ*lutACB* grew in gonococcal base liquid (GCBL) supplemented with glucose, L-lactate, or D-lactate as carbon sources, confirming that LldD and LutACB on their own are both sufficient for L-lactate utilization in this condition (Fig. 1). Δ*lutACB* exhibited a minor but significant growth defect in L-lactate and D-lactate containing media (Fig. 1 B, C, E, F). However, deletion of both *lutACB and lldD* simultaneously (Δ*lutACB* /Δ*lldD)* abolished growth in GCBL supplemented with L-lactate (Fig. 1B, E). As expected, Δ*lutACB* /Δ*lldD* grew equally well in GCBL supplemented with glucose or D-lactate, or on L-lactate when Δ*lutACB* /Δ*lldD* was genetically complemented with *lldD* (Δ*lutACB/lldD+)* confirming that these are the primary active L-LDHs in this condition (Fig. 1).

**Fig 1.**
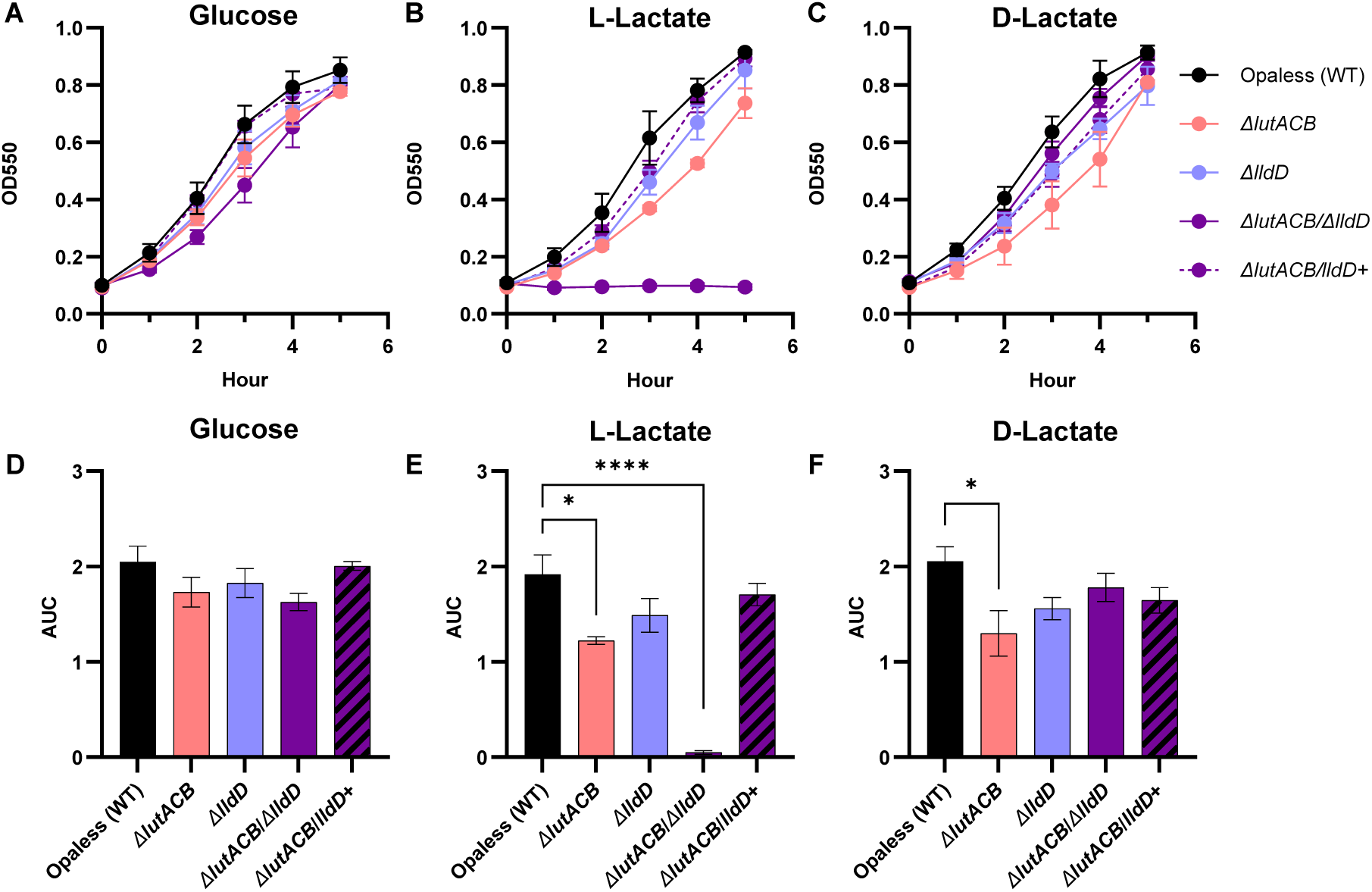
Growth analysis of Gc L-LDH mutants when glucose, L-lactate, or D-lactate is provided as the carbon source. WT Gc and isogenic Δ*lutACB,* Δ*lldD,* Δ*lutACB/lldD,* and Δ*lutACB/lldD+* mutants were cultured in GCBL containing (A and D) 22mM glucose, (B and E) 44mM L-lactate, or (C and F) 44mM D-lactate as the primary carbon source. Growth over 5 hours was monitored by optical density at 550 nm for n = 3-4 biological replicates. (D-F) Area under the curve (AUC) relative to 0h was calculated for each replicate. Significance determined by repeated measures two-way ANOVA with Holm-Šídák’s multiple-comparison test. Symbols and bars represent the mean. Error bars represent SEM. *, *P* < 0.05; ****, *P* < 0.0001.

### LldD, but not LutACB, mediates resistance to superoxide

Exposure of Gc to oxidative stress has been shown to enhance LDH activity in Gc, suggesting that lactate metabolism may contribute to oxidative stress resistance (29). Consistent with these data, mutants in the lactate permease *lctP,* which cannot import lactate, are more susceptible to killing by H_2_O_2_ (25). Thus, we hypothesized that oxidation of L-lactate to pyruvate by LldD and LutACB also supports resistance to oxidative stress in Gc. To test this hypothesis, we examined survival of Gc in the presence of 50 µM paraquat, a superoxide generator, in GCBL (30). Aligning with previous data, we also found that Δ*lctP* was more susceptible to paraquat than WT Gc (Fig. S1) (24). Further, Δ*lldD* Gc exhibited significantly lower survival upon exposure to paraquat compared to WT (Fig 2. A and B). Survival was restored by genetic complementation of *lldD* or by chemical complementation with addition of pyruvate (Fig 2. C and D).

Surprisingly, Δ*lutACB* was entirely dispensable for survival in the presence of paraquat, and Δ*lutACB* /Δ*lldD* was no more sensitive to paraquat than Δ*lldD* (Fig 2. A and B). The defect of Δ*lutACB* /Δ*lldD* survival in the presence of paraquat could be entirely restored by complementation of *lldD* alone (Fig 2. A and B). We hypothesized that *lldD*, but not *lutACB*, is required for resistance to paraquat due to differences in their transcriptional regulation. *lldD* has been found to be significantly upregulated in Gc exposed to H_2_O_2_ by RNAseq (31). *lldD* and *lutACB* have previously been shown to be inversely regulated by iron (27). We hypothesized that expression of *lldD* would therefore be induced and expression of *lutACB* would be dampened in response to incubation with paraquat. We observed that Gc expression of *lldD* was induced by paraquat in a dose and time-dependent manner (Fig. 2E). However, counter to our hypothesis, expression of both *lldD* and *lutA* was induced by paraquat exposure (Fig. 2F). Differences in transcriptional regulation of *lldD* and *lutACB* are therefore not a likely mechanism behind the sensitivity of GC lacking *lldD,* but not *lutACB,* to paraquat.

**Fig 2.**
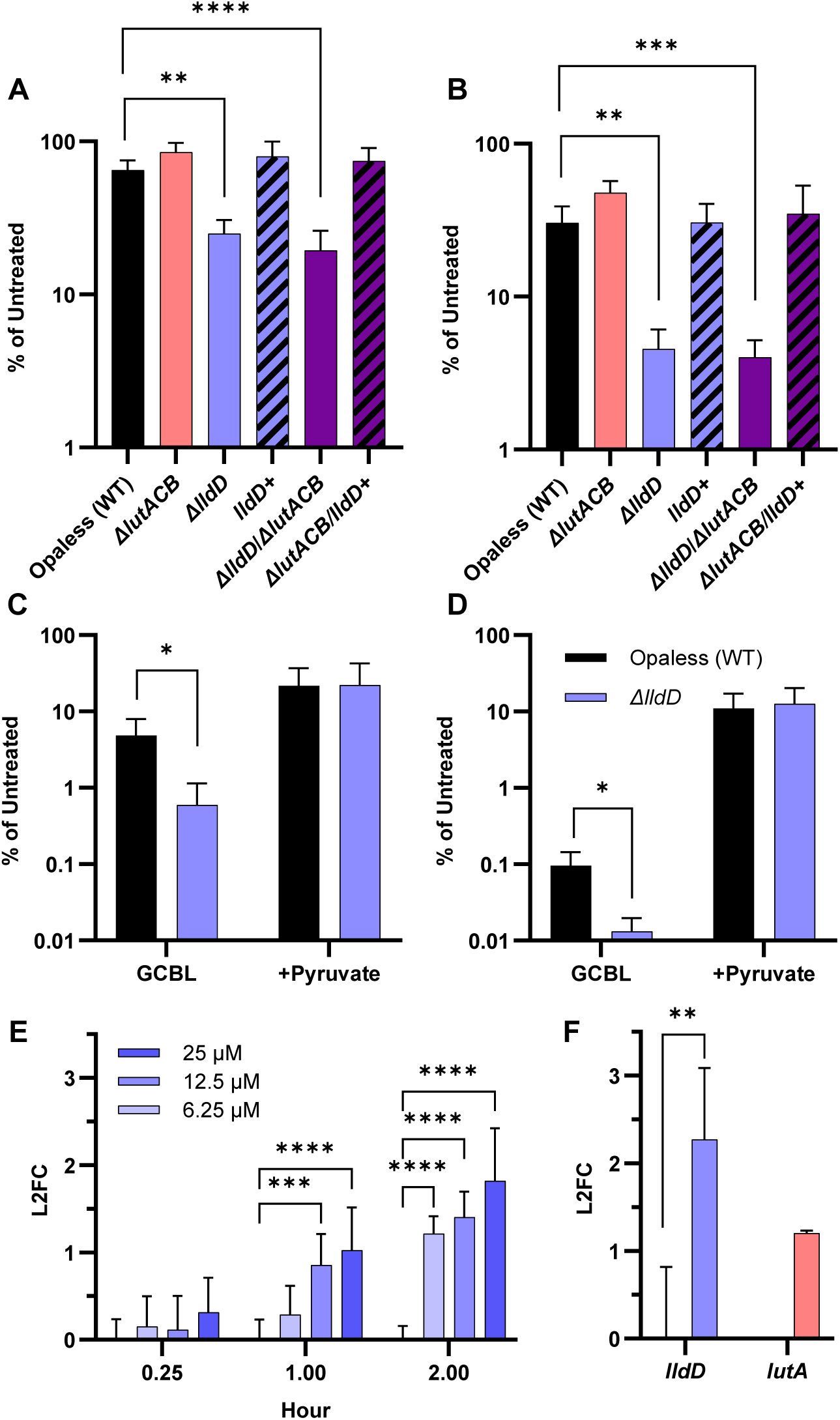
The L-lactate dehydrogenase LldD supports Gc resistance to paraquat derived superoxide. (A-D) WT, Δ*lutACB,* Δ*lldD,* Δ*lldD+,* Δ*lutACB/lldD,* and Δ*lutACB/lldD*+ Gc were exposed to 50 µM paraquat in GCBL, or GCBL supplemented with 44mM pyruvate (shown in C and D). CFU were enumerated after 1 h (A and C) or 2 h (B and D). Bacterial survival is reported relative to the corresponding untreated strain at the same timepoint (set to 100%). Bars represent the mean. Error bars represent SEM. For panels A and B, n = 3-9 biological replicates. Significance determined by mixed effects model with Holm-Šídák’s multiple-comparison test on log-transformed data. For panels C and D, n = 3-4 biological replicates. Significance determined by repeated measures two-way ANOVA with Holm-Šídák’s multiple-comparison test on log-transformed data. (E and F) Gc were exposed to the increasing concentrations of paraquat (up to 25µM in panel E or 50µM in panel F) for the indicated timepoints (up to 2 hours in panel F). RNA was isolated and qRT-PCR was conducted using primers specific for *lldD* (E), *lutA* (E and F) or 16S rRNA (E and F). Gene expression was calculated as 2^-ΔΔCT^ normalized to 16S rRNA, and is expressed relative to untreated controls at each timepoint (log₂-fold change = 0). n = 3 biological replicates. Significance determined by repeated measures two-way ANOVA with Holm-Šídák’s multiple-comparison test. Bars represent the mean. Error bars represent SEM. *, *P* < 0.05; **, *P* < 0.01; ***, *P* < 0.001; ****, *P* < 0.0001.

We next hypothesized that LutACB becomes inactivated by paraquat exposure, rendering it unable to contribute to Gc survival from paraquat. LutACB has an iron-sulfur cluster that is thought to mediate its function; iron-sulfur clusters are known to be oxidized and consequently inactivated by ROS (27, 32). To test this, we measured Gc consumption of L-lactate from media in the presence and absence of paraquat (Fig. S2 A). As expected, we found that WT Gc consumed L-lactate from the media and Δ*lutACB* /Δ*lldD* was unable to consume L-lactate from the media over the course of 2 hours. We observed a partial reduction in L-lactate consumption in both Δ*lldD* and Δ*lutACB.* Strikingly, we observed equivalent L-lactate consumption in Δ*lldD* and Δ*lutACB* in the presence and absence of paraquat, suggesting that both LldD and LutACB are produced and functional in these conditions. We further observed no statistically significant increase in the consumption of lactate upon exposure to paraquat in WT Gc. As confirmation of these results, because L-LDHs are known to convert L-lactate to pyruvate, we also measured pyruvate production by Gc (Fig. S2B). As expected, Δ*lutACB* /Δ*lldD* was unable to produce pyruvate, and production of pyruvate was reduced in Δ*lldD* and Δ*lutACB* compared to WT Gc. A statistically significant, but minor increase in pyruvate production was observed in the presence of paraquat compared to untreated Gc. Together, these results suggest that the survival advantage conferred by LldD is not dependent on lactate consumption or pyruvate production *per se*. Further, although *lutACB* is transcribed and the activity of LutACB is retained during paraquat exposure, its function does not contribute to paraquat resistance. The mechanism of the Δ*lldD-*specific defect in the presence of paraquat is therefore unclear.

### Both *lldD* and *lutACB* deletion mutants are impaired for survival during co-culture with primary human PMNs

PMNs are the primary innate immune cell recruited during Gc infection and are a potent source of ROS through the oxidative burst. Yet, Gc survives in the presence of PMNs (8, 28, 33, 34). Further, PMN derived lactate can serve as a carbon source for Gc (35, 36). We hypothesized that the L-LDHs confer a survival advantage in the presence of PMNs, either due to consumption of PMN-derived lactate or protection from PMN-derived ROS. For these experiments, we used the Opaless Gc strain background, which remains largely extracellular and does not trigger a PMN oxidative burst. PMNs were isolated from whole blood of human donors and infected with Opaless WT, Δ*lldD*, Δ*lutACB, or* Δ*lutACB* /Δ*lldD* in RPMI + 10% FBS to support bacterial growth in the presence of PMNs. We found that Δ*lldD*, Δ*lutACB,* and Δ*lutACB* /Δ*lldD* had significant survival defects in the presence of PMNs at 2 hours post-infection compared to WT (Fig. 3A). This defect could be genetically complemented in Δ*lldD* (Fig. S3). The survival defect in the presence of PMNs was not due to a general growth defect, as Δ*lldD* and Δ*lutACB* cultured in media in the absence of PMNs grew similarly to WT (Fig. 3B). Δ*lutACB* /Δ*lldD* did exhibit a growth defect in media alone, possibly due to a decreased ability to use the small amount of lactate found in the serum component of RPMI +10% FBS, which we measured at ∼ 1mM.

**Fig 3.**
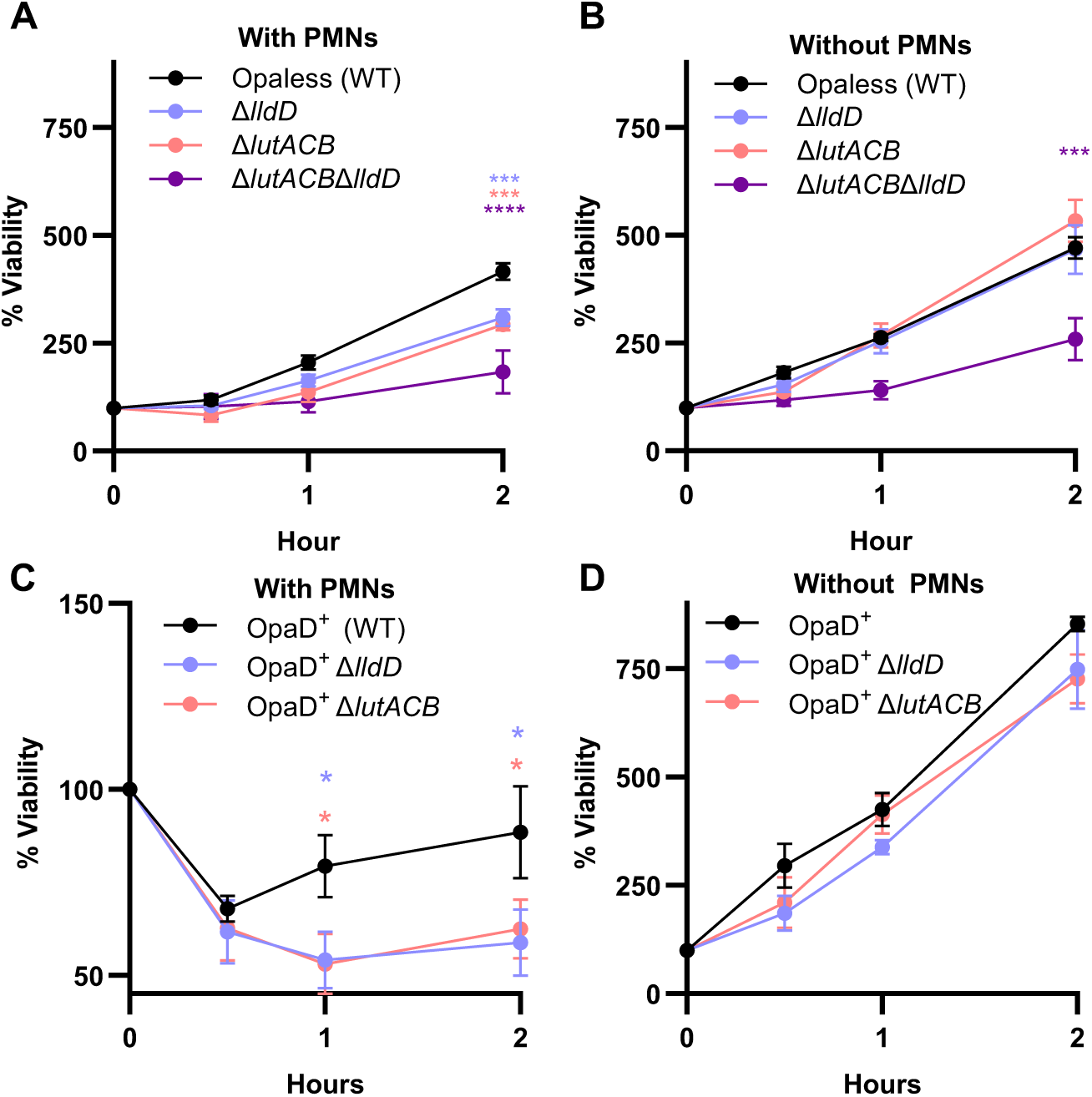
Gc L-lactate dehydrogenases contribute to bacterial survival from PMNs. WT, Δ*lutACB,* Δ*lldD,* and Δ*lutACB/lldD* Gc in the Opaless strain background (A) or an isogenic non-variable OpaD^+^ strain (C and D) were inoculated into IL8-treated PMNs in suspension (A and C), or into media without PMNs (B and D) and incubated for 2 hours. Percent viability was calculated by enumerating Gc CFUs from PMN lysates or media alone controls at 0, 30 mins, 1 hour, or 2 hours post-inoculation, and reporting as % CFU relative to the respective strain at 0 hours. Significance was determined by mixed-effects model with Holm-Šídák’s multiple-comparison test. Symbols indicate the mean and error bars represent SEM. *, *P* < 0.05; **, *P* < 0.01; ***, *P* < 0.001; ****, *P* < 0.0001.

We hypothesized that because LldD confers a survival advantage in the presence of reactive oxygen species, it would also confer a selective survival advantage in the presence of PMN derived ROS. To test this hypothesis, we examined survival of Δ*lldD* and Δ*lutACB* in a constitutively OpaD^+^ expressing Gc strain background, which are rapidly internalized and elicit a robust ROS response within 20 minutes of co-incubation with PMNs (28). However, Δ*lldD*, Δ*lutACB,* and Δ*lutACB* /Δ*lldD* mutants exhibited a comparable magnitude of survival defect, whether in the Opaless or OpaD⁺ background strain (Fig 3. C and D, Fig. S4). Together, this data suggests that the survival defect of L-LDH mutants within PMNs is not dependent on the oxidative burst and may instead be solely due to defects in outgrowth.

### LldD is required for host colonization in a murine model of gonorrhea

The Gc lactate permease was previously reported to promote vaginal colonization in a murine model of gonorrhea (12). As such, we sought to identify if either LldD or LutACB are required for lactate utilization during host infection. To this end, the survival of Δ*lldD* and Δ*lutACB* were compared to WT in mono-infections of the female mouse genital tract. Surprisingly, Δ*lldD* was cleared significantly faster (Fig 4A) and exhibited significantly lower bacterial burdens compared to WT at day 1 and 3 post infection (Fig 4B). This defect was no longer statistically significant at day 5 post-infection, suggesting dependence on *lldD* early in infection. Complementation of *lldD* restored the ability of the mutant to survive within the murine genital tract (Fig. 4). In contrast, *ΔlutACB* colonized the female mouse genital tract equivalently to WT, both in time to clearance and bacterial burden over time (Fig. 4). Together, these data indicate that LldD specifically promotes the survival of Gc *in vivo*, while LutACB is dispensable.

**Fig 4.**
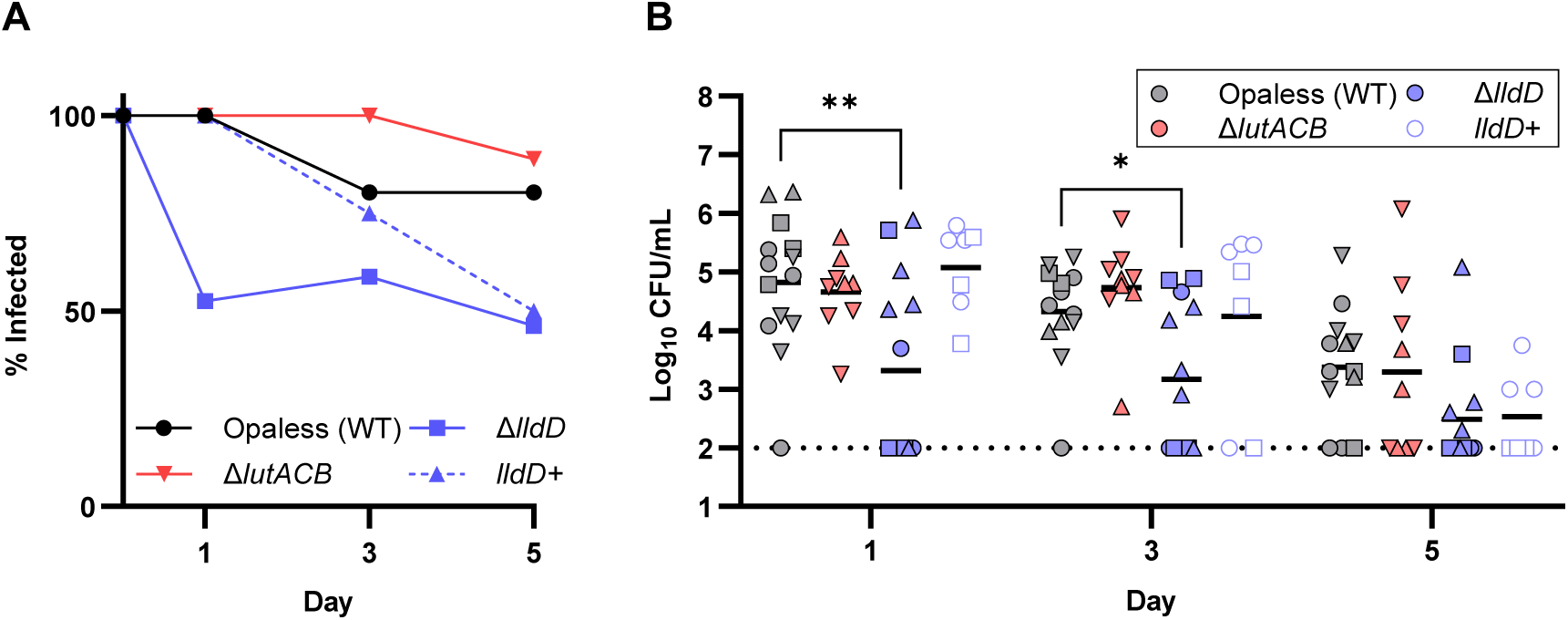
The L-lactate dehydrogenase LldD promotes vaginal colonization in the murine model of gonorrhea. BALB/c mice were infected vaginally with 1*10^6^ CFU of WT, Δ*lutACB,* Δ*lldD,* and *lldD+* Gc in the Opaless strain background. Bacteria were recovered from mice by vaginal swabbing into 1ml of GCBL, and CFU were enumerated at 1, 3, and 5 days post-infection. (A) Percent of mice with at least one CFU of Gc recovered from vaginal swabs compared to the total mice infected at day 0. (B) Each symbol represents data collected from a different mouse. n =7-14. Infection with each strain was repeated at least twice. Shapes represent independent experiments. Black bars indicate the mean. Dotted line represents the limit of detection of the assay. Significance determined by mixed effects model with Holm-Šídák’s multiple-comparison test. *, *P* < 0.05; **, *P* < 0.01.

## Discussion

Lactate is a key metabolite that supports Gc survival *in vivo* and in the presence of PMNs (12, 26, 27). Gc is unusual in encoding four lactate dehydrogenases, two L-lactate specific and two D-lactate specific enzymes (26, 27). Here, we interrogated the two L-lactate dehydrogenases, LldD and LutACB. While it is apparent that either LldD or LutACB are sufficient for growth on L-lactate *in vitro*, our data suggest that LldD and LutACB are not entirely redundant proteins. Here, we identify LldD as a critical enzyme in Gc that supports survival during PMN co-culture, murine colonization, and oxidative stress resistance through differing mechanisms.

Gc is thought to exploit PMN-derived lactate as a nutrient source during infection (35, 36). Previous work, prior to the discovery of *lutACB* in Gc, demonstrated that *lldD* is dispensable for survival during PMN co-culture in Hank’s Buffered Saline Solution (HBSS+), which lacks essential nutrients and prevents bacterial outgrowth (26). However, we found that in a medium that supports bacterial replication (RPMI+10% FBS), both *lldD* and *lutACB* supported survival in the presence of PMNs (Fig. 3). The comparable survival defects observed for Δ*lldD* and Δ*lutACB* across both Opa-expressing and -deficient backgrounds indicate that these L-LDHs promote survival in the presence of PMNs independent of oxidative killing (Fig. S4). These results are consistent with previous reports that the PMN oxidative burst is entirely dispensable for Gc killing (33). Instead, the defects of Δ*lldD* and Δ*lutACB* during PMN co-culture are similar and are likely due to reduced outgrowth ability. This is consistent with previous reports that use of lactate stimulates Gc metabolism and promotes a more rapid entry into log-phase growth (11). It is unclear why LldD and LutACB are able to individually support growth in media but not during PMN co-culture, though decreased overall fitness of Gc in stressed conditions may play a role.

Lactate utilization is also thought to contribute to Gc pathogenesis *in vivo*. The gonococcal lactate permease was previously shown to promote colonization of the murine genital tract (12). Our results expand upon this finding, implicating LldD, but not LutACB, as conferring a fitness advantage *in vivo* during murine colonization (Fig. 4). The exact mechanism of this advantage is unclear. Although Δ*lldD* has a survival defect in the presence of PMNs *in vitro*, PMNs likely do not mediate the survival defect *in vivo*. During Gc infection of BALB/c mice, PMNs are typically recruited by day 5 post-infection (37). In our conditions, we observed no PMN recruitment at days 1 and 3 post-infection. It is therefore unlikely that PMNs contribute significantly to L-lactate levels within the genital tract in mice, particularly at the early timepoints for which a survival defect was observed for Δ*lldD in vivo* (Fig. 4). However, lactate is also produced by host epithelial cells and residual microbiota within the genital tract that persist during the antibiotic treatment used to facilitate Gc colonization (26, 27). Lactate is therefore abundant within the human and mouse genital tracts even in the absence of infection (12–17). In this context, LldD appears to be the primary L-LDH that Gc uses to capitalize on available L-lactate early in infection.

Lactate utilization has previously been linked to oxidative stress resistance in Gc. Prior work determined that Δ*lctP* Gc are more susceptible to killing by ROS (24). Our results support this conclusion and expand upon it to reveal that the L-LDH LldD confers oxidative stress resistance in Gc. Many mediators of oxidative stress resistance are required for Gc fitness *in vivo*, including *kat* (catalase), *msrAB* (methionine sulfide reductase), and *mntC* (38). We speculate that the defects of both Δ*lctP* and Δ*lldD in vivo* are due to increased susceptibility to oxidative stress, and that L-lactate catabolism is integrated into the broader oxidative stress response *in vivo*. It is important for future work to clarify if the *in vivo* fitness defects of Δ*lldD* and Δ*lctP* are due specifically to an inability to consume L-lactate as a carbon source, corresponding defects in ROS resistance, or some other undetermined factor.

We found that both *lldD* and *lutA* are induced in response to paraquat (Fig. 2 E and F). This aligns with previous data that *lldD* is upregulated in response to ROS such as H_2_O_2_ (31). However, *lldD* and *lutACB* are known to be differentially regulated (27). In Gc, *lutACB* contains a canonical ferric uptake regulator (Fur) binding site and is activated under iron-replete conditions, whereas *lldD* expression increases under iron-limited conditions (27). This regulation aligns with the catalytic mechanism of LutACB, which contains several iron-sulfur clusters within the enzyme complex (32). In contrast, LldD is a flavoprotein that does not require iron (32). Iron limitation would be expected to favor iron-sparing enzymes. Both PMN co-culture and murine infection are iron limited due to the presence of iron-binding proteins such as bovine transferrin found in FBS during PMN co-culture *in vitro* and murine transferrin, hemoglobin, and lactoferrin *in vivo* which may favor *lldD* expression (39–41). In contrast, during culture in the iron-replete medium GCBL *in vitro*, we would expect that *lutACB* is upregulated and *lldD* expression is repressed. Iron availability is therefore likely a major determinant of which L-LDH dominates in a given environment. Further, reduction of iron transport in Gc can prevent ROS-mediated killing, by minimizing the availability of redox-active labile iron, explicitly linking lactate utilization, metal homeostasis, and oxidative stress resistance (42).

Interestingly, *lutACB* was previously linked to resistance from H_2_O_2_ in *N. meningitidis* (NMB1436–38), prior to the identification of its function as an L-LDH (43). Although Δ*lutACB* did not exhibit a survival defect in the presence of paraquat in Gc, we did observe induction of *lutA* by paraquat along with *lldD*, suggesting that both L-lactate dehydrogenases are expressed concurrently in iron-replete, ROS-rich environments. We also observed that both Δ*lldD* and Δ*lutACB* consumed similar amounts of lactate upon paraquat exposure (Fig. S2A), further supporting that both L-LDHs are active in our conditions. One caveat is that the decreased viability of Δ*lldD* and Δ*lutACB /lldD* upon exposure to paraquat may confound bulk measurements of L-lactate consumption from the medium. Approaches that can assess L-lactate consumption rate normalized to viable cell density or directly measure conversion of L-lactate to pyruvate may be more sensitive for detecting subtle differences in LDH activity. Regardless, our data indicate that LldD and LutACB promote survival under oxidative stress through independent roles beyond straightforward differences in transcriptional regulation or gross catalytic activity.

Alternative explanations for the sensitivity of lactate utilization mutants to ROS have been proposed. It has been speculated that pyruvate may act directly as an antioxidant to protect Gc against the activity of ROS (24). However, the differences in ROS sensitivity of Δ*lldD* and Δ*lutACB* in the presence of paraquat despite similar production of pyruvate (Fig. S2B) suggest this may not be the case. Instead, the sensitivity of Δ*lldD* to ROS may be tied to other downstream consequences of lactate catabolism. In support of this hypothesis, chemical complementation with pyruvate rescued the defect of Δ*lldD* compared to WT (Fig. 2 C and D), suggesting that catabolism of L-lactate into pyruvate, rather than the action of the *lldD* itself, may support ROS resistance. There is some evidence that this is mediated by transcriptional changes as there is significant overlap between the lactate, iron, and H_2_O_2_ responsive regulons in Gc (25). Consistent with this data, L-lactate suppresses transcription of genes related to iron transport (25). Together, these observations support a model in which L-lactate catabolism supports ROS resistance, in part, by regulation of iron uptake to support redox homeostasis. Whether pyruvate can elicit similar transcriptional changes, and whether LldD and LutACB differentially contribute to these changes remains to be determined.

LldD and LutACB have distinct biochemical properties that may dictate their roles in oxidative stress resistance and survival *in vivo*. A thorough review of the biochemical properties of bacterial LDHs was recently published (32). Briefly, the LldD-type L-LDHs are single-subunit, FMN-dependent proteins that transfer electrons directly from L-lactate to the quinone pool (44). Although LldD-type LDHs are highly specific for L-lactate, some can also use alternative substrates such as α-hydroxybutyrate (44). In contrast, LutACB-type L-LDHs are tripartite, flavin-independent enzymes (45, 46). LutACB-type L-LDHs are also specific for L-lactate, however, overexpression of these proteins can allow for use of D-lactate as a growth substrate in *E. coli* (45). LutACB-type LDHs are thought to relay electrons from lactate oxidation by LutC across iron-sulfur clusters within LutA and B to quinones(47). These structural differences suggest several hypotheses for why LldD and LutACB are not entirely redundant in Gc: differences in (i) affinity and specificity for L-lactate, (ii) efficiency of electron transfer to the respiratory chain, (iii) vulnerability to oxidative inactivation, and (iv) availability of essential co-factors. These possibilities are not mutually exclusive and in-depth biochemical studies will be required to distinguish between them.

Together, our results call for a reassessment of prior conclusions regarding lactate utilization in Gc and, by extension, other *Neisseria* species. Earlier studies linking lactate permease mutants to virulence defects broadly attributed these effects to impaired lactate utilization; however, we now know that the lactate permease also functions as a pyruvate transporter (25). Our data instead demonstrate that individual lactate dehydrogenases exhibit distinct contributions to growth and infectivity. In conclusion, we expand upon previous work implicating lactate utilization in Gc virulence and reveal that Gc L-LDHs serve both overlapping and distinct functions across multiple infection contexts, including oxidative stress, PMN challenge, and colonization *in vivo*. Given the conservation of L-LDHs across *Neisseria* species (32), these enzymes likely contribute to niche-specific physiology and adaptation, warranting renewed investigation into their roles in supporting bacterial survival and pathogenesis across diverse infection sites.

## Materials and Methods

### Ethics Statement

Human subjects provided informed consent in accordance with protocol #202408109 or #13909, approved by the University of Iowa Carver College of Medicine Institutional Review Board and the University of Virginia Institutional Review Board for Health Sciences Research, respectively. The murine model of gonorrhea was performed in accordance with protocol # 4062608, approved by the University of Iowa Carver College of Medicine Institutional Animal Care and Use Committee.

### Bacterial Strains and Growth Conditions

Opaless is a non-variable Opa-deficient derivative of Gc strain FA1090, which constitutively expresses the 1-81-S2 pilin variant (28). Strain 1291 Δ*lldD* and 1291 Δ*lutACB* were received from Dr. Jennifer Edwards (26). Strain F62 Δ*lctP* was obtained from Dr. Ann Jerse (12). DNA containing pCTS32::*lldD* for genetic complementation of Δ*lldD* was received from Dr. Alastair McEwan (27). Genomic DNA was used for transformation of mutated genes into the Opaless strain over three successive backcrosses. Genomic DNA from metabolic mutants in the Opaless Gc background were used to transform the isogenic, non-variable OpaD^+^ Gc background (28). Mutants were verified by PCR or whole genome sequencing.

Gc were grown at 37°C with 5% CO_2_ on Gonococcal Medium Base (GCB, Difco) plus Kellogg’s supplements (48) or on Chocolate Agar (Hardy Diagnostics). Logarithmic phase Gc were prepared by growing Gc in liquid medium (GCBL) plus Kellogg’s supplements for successive rounds of dilution with enrichment for piliation, as previously described (49).

Kanamycin (50 µg/ml) was used for selection of *lldD* mutants. Chloramphenicol (0.5 µg/ml) was used for selection of *lutACB* and *lctP* mutants. Spectinomycin (80 µg/ml) was used for selection of *lldD*+.

### Growth Curves

Gc was grown in GCBL containing 22mM glucose (found in Kellog’s supplements), or carbon-equivalents matched GCBL in which glucose was replaced with 44mM L-lactate or D-lactate. In brief, mid-logarithmic phase Gc were pelleted and resuspended in 15 ml conical tubes at ∼5*10^7^ CFU/ml in 6mls of the desired media. Growth was monitored every hour for 5 total hours by OD550.

### Paraquat survival Assays

Mid-logarithmic phase Gc were diluted to ∼5*10^7^ CFU/ml in GCBL, or in GCBL supplemented with 44mM L-lactate, D-lactate, or pyruvate for chemical complementation studies. Gc was then treated with the indicated concentration of paraquat or left untreated. CFUs were enumerated at the indicated timepoints post-treatment relative to the untreated control (100%).

### RNA extraction, cDNA synthesis, and qRT-PCR

Bacteria (>1ml) were harvested at the indicated timepoints from paraquat survival assays, pelleted, and resuspended in RNAlater stabilization solution, and stored at 4°C until processed for RNA extraction. Samples were lysed according to the “Sample Lysis of Bacteria or Yeast” protocol before proceeding with the protocol for “Total RNA Isolation and Purification” Using the Monarch Spin RNA Isolation Kit (Mini) (New England Biolabs;#T2110). cDNA synthesis was then performed from the RNA product using the High-Capacity cDNA Reverse Transcription Kit (Applied Biosystems). qPCR was conducted using a Thermo Fisher Scientific QuantStudio3 instrument, *Power* SYBR green PCR master mix (Applied Biosystems 4367659), and previously described qRT-PCR primers for *lldD*, *lutA*, and a *16S* control (27). Data were normalized to a 16S rRNA reference control. Data are expressed as expression (2^-ΔΔCT^) relative to the untreated control condition at each timepoint.

### Metabolite quantification

Gc were harvested at 2 hours post treatment with paraquat, pelleted, and supernatants were removed and stored at −80°C until processed for lactate and pyruvate concentrations. L-lactate and pyruvate concentrations were measured using L-lactate assay kit (Abcam) and pyruvate assay kit (Abcam) and calculated using a standard curve according to manufacturer instructions.

### Gc-PMN Co-culture

Primary human PMNs were isolated from venous blood of subjects by dextran sedimentation, Ficoll-Paque™ separation, and osmotic red blood cell lysis as previously described and utilized within 2 hours of collection (50). As previously described, PMNs in suspension (1*10^6^ PMNs/ml) were treated with 10 nM human IL-8 (R&D Biosystems) in Roswell Park Memorial Institute 1640 medium (RPMI-1640 without L-glutamine; Cytiva HyClone) supplemented with 10% heat-inactivated fetal bovine serum (FBS) (35). Gc was added to each conical at a multiplicity of infection of 10 (1*10^7^ CFU/ml) or to a culture containing media without PMNs and incubated at 37°C. CFU counts were enumerated from at the specified timepoints and expressed relative to CFU at 0 hours (100%).

### Murine genital infection

Infection of female β-estradiol treated BALB/c mice was performed as previously described (51). Briefly, female BALB/c mice at 4-6 weeks old were ordered from Charles River and allowed to acclimate for 10 days in a female only room of the animal facility prior to beginning the experiment. Mice were housed with autoclaved food, water, bedding and cage components for the duration of the experiment. Mice in diestrus or anestrus stage were given β-estradiol (5mg/ml) in 100 μl sesame oil subcutaneously, and streptomycin (24 mg/ml) and vancomycin (4mg/ml) intraperitoneally (Day −2: 200 μl; Day −1: 200 μl, Day 0: 150 μl; Day 1: 150 μl) to knock down commensal overgrowth under the influence of estradiol. Trimethoprim sulfate (0.4g/L; Day −2, 0, 2, and 4) and streptomycin (5g/L; Day 2 and 4) were added to the drinking water to further limit commensal outgrowth. Mice were infected with 1*10^6^ CFU in 20 μl PBS. Bacterial burden was enumerated on alternate days by vaginal swabbing into 1ml of GCBL for 5 days following inoculation. Mice with overgrowth of gram-negative rods were removed from the study and subsequent calculations. Percent infected was calculated as number of mice with any Gc colonies identified above the limit of detection out of the total mice infected.

## Acknowledgements

This work was supported in part by NIH T32AI007496, NIH R01AI127793-07W2, and startup funds from the University of Iowa Carver College of Medicine to ADP. AKC was supported by NIH R01AI097312, NIH R01AI127793, R21AI161302, and the Harrison Distinguished Teaching Professorship.

## Supplemental Material

**Fig S1:**
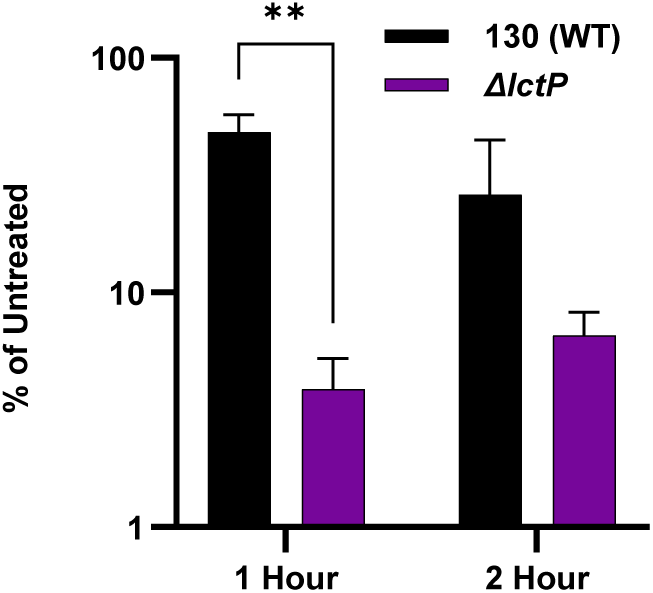
Lactate permease is required for full resistance of Gc to paraquat derived superoxide. WT and Δ*lctP* Gc were exposed to 50 µM paraquat in GCBL. CFU were enumerated post-exposure at the indicated timepoint. Bacterial burden is reported relative to the corresponding untreated strain at that timepoint (100%). Bars represent the mean. Error bars represent SEM. (A and B) n = 3-5 biological replicates. WT replicates represent a subset of data reported in Fig 2A and B. Significance determined by mixed effects model with Holm-Šídák’s multiple-comparison test on log-transformed data. ****, *P* < 0.0001.

**Fig S2:**
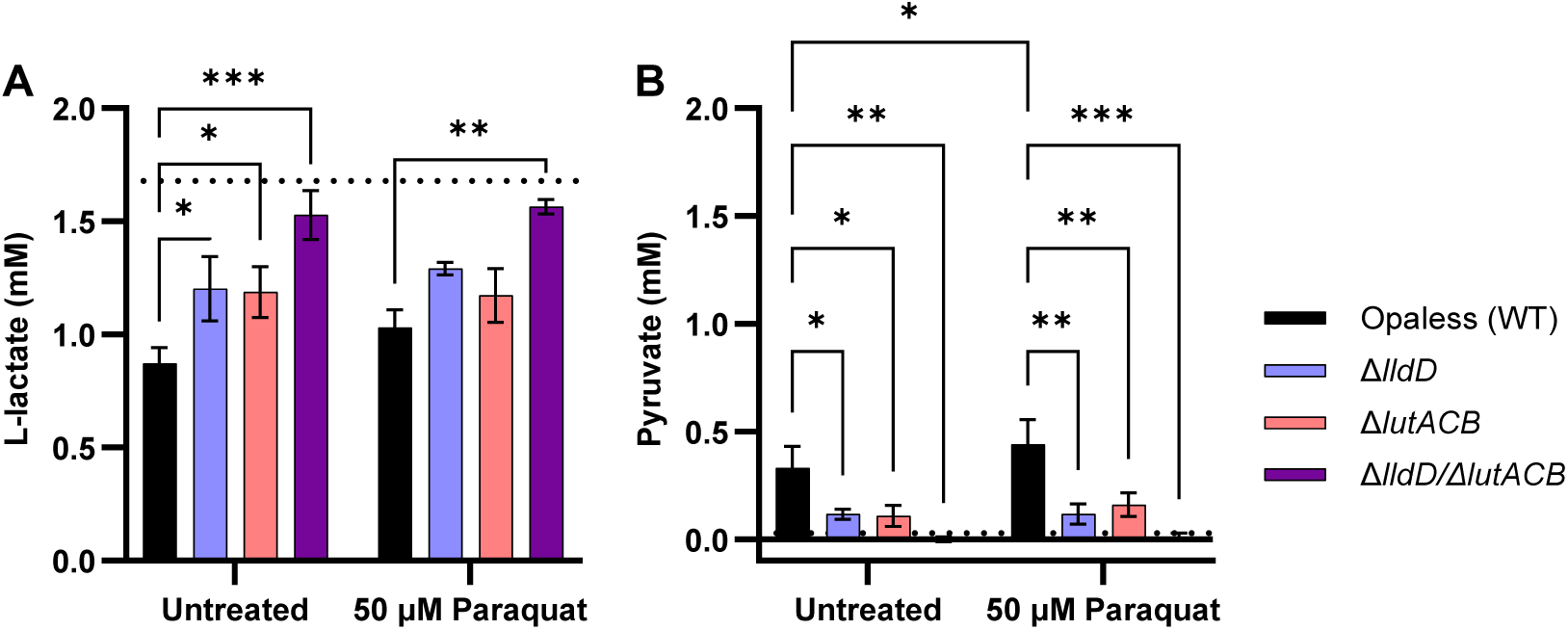
Consumption of L-lactate and production of pyruvate is largely unchanged with exposure to paraquat. WT Gc, Δ*lutACB,* Δ*lldD,* and Δ*lutACB/lldD* were exposed to 50 µM paraquat in GCBL. Bacteria were collected, pelleted, and supernatants were retained for measurement of (A) L-lactate and (B) pyruvate. Bars represent the mean. Error bars represent SEM. Dotted line indicates baseline metabolite concentration in media at time 0. n = 3 biological replicates. Significance determined by mixed effects model with Holm-Šídák’s multiple-comparison test. *, *P* < 0.05; **, *P* < 0.01; ***, *P* < 0.001; ****.

**Fig S3:**
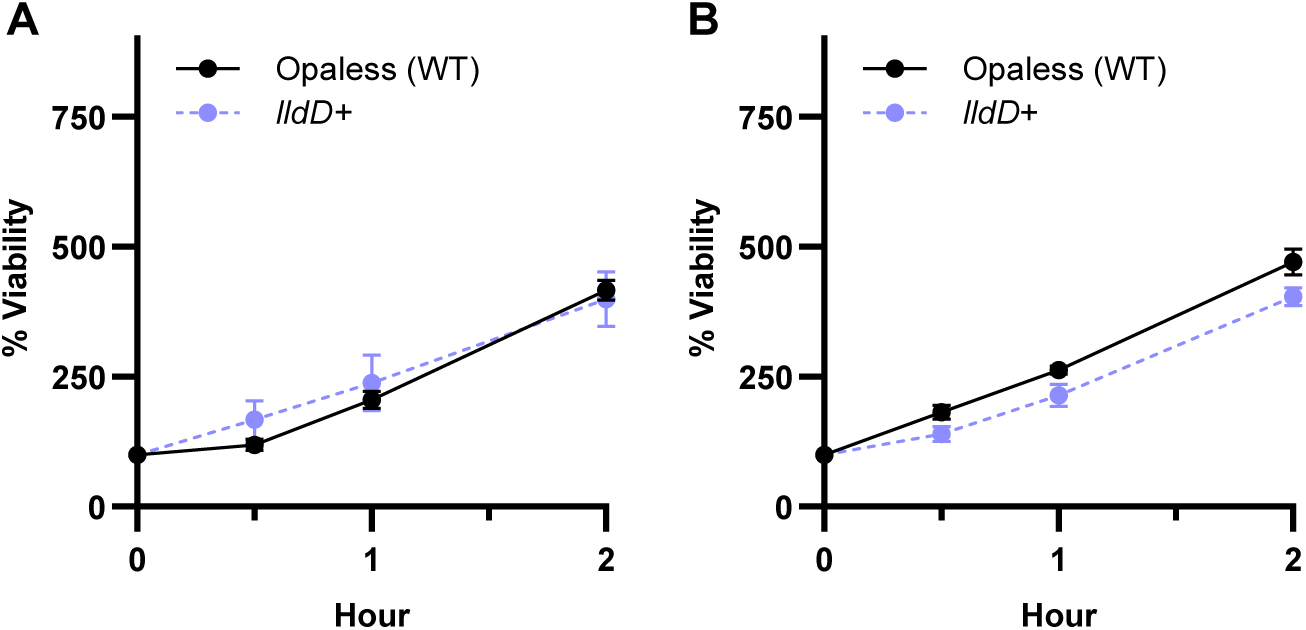
Genetic complementation restores the survival defect of *lldD*+ in the presence of PMNs. WT and *lldD+* Gc in the Opaless strain background were inoculated into IL8-treated PMNs in suspension (A), or into media without PMNs (B) and incubated for 2 hours. Percent viability was calculated by enumerating Gc CFUs from PMN lysates or media alone controls at 0, 30 mins, 1 hour, or 2 hours post-inoculation, and reporting as % CFU relative to the respective strain at 0 hours. Significance was determined by mixed effects model with Holm-Šídák’s multiple-comparison test. Symbols indicate the mean and error bars represent SEM. n = 5. WT replicates represent a subset of data reported in Fig 3A and B.

**Fig S4.**
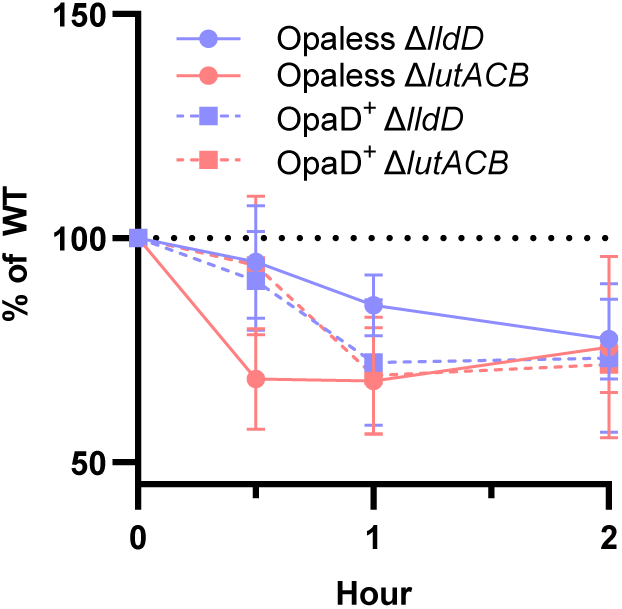
Gc L-lactate dehydrogenases contribute to bacterial survival from PMNs. WT, Δ*lutACB,* and Δ*lldD* Gc in the Opaless strain background or an isogenic non-variable OpaD^+^ strain were inoculated into IL8-treated PMNs in suspension and incubated for 2 hours. Percent of WT survival was calculated by enumerating CFUs from PMN lysates or media alone controls at 0, 30 mins, 1 hour, or 2 hours post-inoculation, and reporting as % CFU relative to the respective strain at 0 hours, normalized to the WT at that timepoint. Significance was determined by mixed effects model with Holm-Šídák’s multiple-comparison test. No significant difference between Opaless and OpaD^+^ strains was detected at any timepoint.

## References

1. A. Curry, T. Williams, M. L. Penny, Pelvic Inflammatory Disease: Diagnosis, Management, and Prevention. American family physician 100 (2019).

2. D. Tsevat, H. Wiesenfeld, C. Parks, J. Peipert, Sexually transmitted diseases and infertility. American journal of obstetrics and gynecology 216 (2017).

3. A. Lovett, J. Duncan, Human Immune Responses and the Natural History of Neisseria gonorrhoeae Infection. Frontiers in immunology 9 (2019).

4. A. D. Potter, A. K. Criss, Dinner date: Neisseria gonorrhoeae central carbon metabolism and pathogenesis. Emerging topics in life sciences 10.1042/ETLS20220111 (2023).

5. C. N. Cornelissen, Subversion of nutritional immunity by the pathogenic *Neisseriae*. Pathog Dis 76, ftx112 (2018).

6. S. J. Quillin, H. S. Seifert, *Neisseria gonorrhoeae* host adaptation and pathogenesis. Nature reviews. Microbiology 16, 226–240 (2018).

7. I. K. Liyayi, A. L. Forehand, J. C. Ray, A. K. Criss, Metal piracy by Neisseria gonorrhoeae to overcome human nutritional immunity. PLoS pathogens 19 (2023).

8. A. Palmer, A. K. Criss, Gonococcal Defenses against Antimicrobial Activities of Neutrophils. Trends Microbiol 26, 1022–1034 (2018).

9. W. Song, Q. Yu, L. C. Wang, D. C. Stein, Adaptation of Neisseria gonorrhoeae to the Female Reproductive Tract. Microbiol Insights 13, 1178636120947077 (2020).

10. H. Smith, C. M. Tang, R. M. Exley, Effect of host lactate on gonococci and meningococci: new concepts on the role of metabolites in pathogenicity. Infect Immun 75, 4190–4198 (2007).

11. H. Smith, E. A. Yates, J. A. Cole, N. J. Parsons, Lactate stimulation of gonococcal metabolism in media containing glucose: mechanism, impact on pathogenicity, and wider implications for other pathogens. Infect Immun 69, 6565–6572 (2001).

12. R. M. Exley et al., Lactate acquisition promotes successful colonization of the murine genital tract by *Neisseria gonorrhoeae*. Infect Immun 75, 1318–1324 (2007).

13. S. Al-Mushrif, A. Eley, B. Jones, Inhibition of chemotaxis by organic acids from anaerobes may prevent a purulent response in bacterial vaginosis. Journal of medical microbiology 49 (2000).

14. G. R. Huggins, G. Preti, Volatile constituents of human vaginal secretions. Am J Obstet Gynecol 126, 129–136 (1976).

15. G. Preti, G. R. Huggins, G. D. Silverberg, Alterations in the organic compounds of vaginal secretions caused by sexual arousal. Fertil Steril 32, 47–54 (1979).

16. D. H. Owen, D. F. Katz, A vaginal fluid simulant. Contraception 59, 91–95 (1999).

17. D. E. O’Hanlon, T. R. Moench, R. A. Cone, Vaginal pH and microbicidal lactic acid when lactobacilli dominate the microbiota. PLoS One 8, e80074 (2013).

18. E. Boskey, R. Cone, K. Whaley, T. Moench, Origins of vaginal acidity: high D/L lactate ratio is consistent with bacteria being the primary source. Human reproduction (Oxford, England) 16 (2001).

19. S. Morse, Neisseria gonorrhoeae: physiology and metabolism. Sexually transmitted diseases 6 (1979).

20. E. Yates et al., In a medium containing glucose, lactate carbon is incorporated by gonococci predominantly into fatty acids and glucose carbon incorporation is increased: implications regarding lactate stimulation of metabolism. International journal of medical microbiology: IJMM 290 (2000).

21. L. Gao, N. J. Parsons, A. Curry, J. A. Cole, H. Smith, Lactate causes changes in gonococci including increased lipopolysaccharide synthesis during short-term incubation in media containing glucose. FEMS Microbiol Lett 169 (1998).

22. T. Regan, A. Watts, H. Smith, J. Cole, Regulation of the lipopolysaccharide-specific sialyltransferase activity of gonococci by the growth state of the bacteria, but not by carbon source, catabolite repression or oxygen supply. Antonie Van Leeuwenhoek 75, 369–379 (1999).

23. A. J. Cardenas et al., Neisseria gonorrhoeae scavenges host sialic acid for Siglec-mediated, complement-independent suppression of neutrophil activation. mBio 15, e0011924 (2024).

24. J. C. Ayala, W. M. Shafer, Transcriptional regulation of a gonococcal gene encoding a virulence factor (L-lactate permease). PLoS pathogens 15 (2019).

25. J. C. Ayala, J. T. Balthazar, W. M. Shafer, Transcriptional responses of Neisseria gonorrhoeae to glucose and lactate: implications for resistance to oxidative damage and biofilm formation. mBio 15, e0176124 (2024).

26. J. M. Atack et al., A Role for Lactate Dehydrogenases in the Survival of *Neisseria gonorrhoeae* in Human Polymorphonuclear Leukocytes and Cervical Epithelial Cells. J Infect Dis 210, 1311–1318 (2014).

27. N. H. Chen et al., Two Distinct L-Lactate Dehydrogenases Play a Role in the Survival of *Neisseria gonorrhoeae* in Cervical Epithelial Cells. J Infect Dis 221, 449–453 (2020).

28. L. M. Ball, A. K. Criss, Constitutively Opa-expressing and Opa-deficient *Neisseria gonorrhoeae* strains differentially stimulate and survive exposure to human neutrophils. J Bacteriol 195, 2982–2990 (2013).

29. H. S. Fu, D. J. Hassett, M. S. Cohen, Oxidant stress in Neisseria gonorrhoeae: adaptation and effects on L-(+)-lactate dehydrogenase activity. Infection and immunity 57 (1989).

30. H. M. Hassan, I. Fridovich, Superoxide radical and the oxygen enhancement of the toxicity of paraquat in Escherichia coli. J Biol Chem 253, 8143–8148 (1978).

31. S. Quillin, A. Hockenberry, M. Jewett, H. Seifert, Neisseria gonorrhoeae Exposed to Sublethal Levels of Hydrogen Peroxide Mounts a Complex Transcriptional Response. mSystems 3 (2018).

32. A. G. McEwan, J. Hosmer, U. Kappler, Molecular and cellular biology of bacterial lactate metabolism. Adv Microb Physiol 87, 299–355 (2025).

33. A. K. Criss, B. Z. Katz, H. S. Seifert, Resistance of Neisseria gonorrhoeae to non-oxidative killing by adherent human polymorphonuclear leucocytes. Cell Microbiol 11, 1074–1087 (2009).

34. A. K. Criss, H. S. Seifert, A bacterial siren song: intimate interactions between Neisseria and neutrophils. Nature reviews. Microbiology 10, 178–190 (2012).

35. A. D. Potter, C. M. Baiocco, J. A. Papin, A. K. Criss, Transcriptome-guided metabolic network analysis reveals rearrangements of carbon flux distribution in Neisseria gonorrhoeae during neutrophil co-culture. mSystems 10.1128/msystems.01265-22 (2023).

36. B. E. Britigan, D. Klapper, T. Svendsen, M. S. Cohen, Phagocyte-derived lactate stimulates oxygen consumption by *Neisseria gonorrhoeae*. An unrecognized aspect of the oxygen metabolism of phagocytosis. J Clin Invest 81, 318–324 (1988).

37. M. Packiam, S. J. Veit, D. J. Anderson, R. R. Ingalls, A. E. Jerse, Mouse strain-dependent differences in susceptibility to Neisseria gonorrhoeae infection and induction of innate immune responses. Infect Immun 78, 433–440 (2010).

38. H. Wu, A. A. Soler-Garcia, A. E. Jerse, A strain-specific catalase mutation and mutation of the metal-binding transporter gene mntC attenuate Neisseria gonorrhoeae in vivo but not by increasing susceptibility to oxidative killing by phagocytes. Infect Immun 77, 1091–1102 (2009).

39. A. E. Jerse et al., Growth of Neisseria gonorrhoeae in the female mouse genital tract does not require the gonococcal transferrin or hemoglobin receptors and may be enhanced by commensal lactobacilli. Infect Immun 70, 2549–2558 (2002).

40. B. C. Lee, A. B. Schryvers, Specificity of the lactoferrin and transferrin receptors in Neisseria gonorrhoeae. Mol Microbiol 2, 827–829 (1988).

41. T. Hagen, C. Cornelissen, Neisseria gonorrhoeae requires expression of TonB and the putative transporter TdfF to replicate within cervical epithelial cells. Molecular microbiology 62 (2006).

42. S. Varghese, A. Wu, S. Park, K. R. Imlay, J. A. Imlay, Submicromolar hydrogen peroxide disrupts the ability of Fur protein to control free-iron levels in Escherichia coli. Mol Microbiol 64, 822–830 (2007).

43. R. Grifantini et al., Characterization of a novel Neisseria meningitidis Fur and iron-regulated operon required for protection from oxidative stress: utility of DNA microarray in the assignment of the biological role of hypothetical genes. Mol Microbiol 54, 962–979 (2004).

44. M. Futai, H. Kimura, Inducible membrane-bound L-lactate dehydrogenase from Escherichia coli. Purification and properties. J Biol Chem 252, 5820–5827 (1977).

45. G. E. Pinchuk et al., Genomic reconstruction of Shewanella oneidensis MR-1 metabolism reveals a previously uncharacterized machinery for lactate utilization. Proc Natl Acad Sci U S A 106, 2874–2879 (2009).

46. Y. Chai, R. Kolter, R. Losick, A widely conserved gene cluster required for lactate utilization in Bacillus subtilis and its involvement in biofilm formation. J Bacteriol 191, 2423–2430 (2009).

47. W. C. Hwang et al., LUD, a new protein domain associated with lactate utilization. BMC Bioinformatics 14, 341 (2013).

48. D. S. Kellogg, Jr., W. L. Peacock, Jr., W. E. Deacon, L. Brown, C. I. Pirkle, *Neisseria gonorrhoeae* I: Virulence Genetically Linked to Clonal Variation. J Bacteriol 85, 1274–1279 (1963).

49. A. K. Criss, H. S. Seifert, *Neisseria gonorrhoeae* suppresses the oxidative burst of human polymorphonuclear leukocytes. Cell Microbiol 10, 2257–2270 (2008).

50. E. A. Stohl, A. K. Criss, H. S. Seifert, The transcriptome response of Neisseria gonorrhoeae to hydrogen peroxide reveals genes with previously uncharacterized roles in oxidative damage protection. Molecular microbiology 58 (2005).

51. E. L. Raterman, A. E. Jerse, Female Mouse Model of Neisseria gonorrhoeae Infection. Methods in molecular biology (Clifton, N.J.) 1997 (2019).

